# Autoreactive T cells preferentially drive differentiation of non-responsive memory B cells at the expense of germinal center maintenance

**DOI:** 10.1101/287789

**Authors:** Rajiv W Jain, Kate A Parham, Yodit Tesfagiorgis, Heather C Craig, Emiliano Romanchik, Steven M Kerfoot

## Abstract

B cell fate decisions within a germinal center (GC) are critical to determining the outcome of the immune response to a given antigen. Here, we characterize GC kinetics and B cell fate choices in a response to the autoantigen myelin oligodendrocyte glycoprotein (MOG), and compare them the response to a standard model foreign antigen (NP-haptenated ovalbumin, NPOVA). Both antigens generated productive primary responses, as evidenced by GC development, circulating antigen-specific antibodies, and differentiation of memory B cells. However, in the MOG response the status of the cognate T cell partner drove preferential B cell differentiation to a memory phenotype at the expense of GC maintenance, resulting in a truncated GC. Reduced plasma cell differentiation was largely independent of T cell influence. Interestingly, memory B cells formed in the MOG GC were unresponsive to secondary challenge and this could not be overcome with T cell help.

## Introduction

Tailoring the immune response to a given antigen is a crucial function of the immune system, as the quality and nature of the response impacts the success of pathogen clearance as well as subsequent long-lived immunity. This is further complicated in cases where the response directly targets or cross reacts with a self-antigen. Nearly all immune responses incorporate both B and T cell recognition of the antigen, and collaboration between B and T cells specific for said antigen produces a germinal center (GC) response (Shlomchik and Weisel, 2012; Victora and Nussenzweig, 2012; Vinuesa et al., 2016). Throughout the GC response, B cell survival, proliferation, and differentiation to either antibody-producing plasma cells or memory B cells is dependent upon, and informed by, direct interactions with T cells specific for the same antigen (cognate interactions) (Mesin et al., 2016). However, the signals that drive differential fate choices made by B cells responding to different antigens and how they are influenced by features of the antigen itself are not well understood.

Interactions with cognate T cells are critical during two distinct phases of the developing B cell response. The first phase occurs shortly after exposure to a new antigen, but prior to GC formation. During this phase, cognate B/T interactions are essential to initiate antigen-stimulated B cell proliferation and also to drive B cell differentiation along three distinct pathways; short-lived plasmablasts that produce low affinity, largely IgM antibodies; early (mostly) IgM memory B cells; and GC B cells that reenter the follicle to initiate a new GC (Corcoran and Tarlinton, 2016). The second phase is within the mature GC itself. GC B cells undergo clonal expansion and somatic hypermutation largely within the dark zone (DZ), before migrating to the light zone (LZ) to compete for survival signals supplied through interactions with specialized cognate T follicular helper (Tfh) cells (Mesin et al., 2016). Evidence also suggests that Tfh cells provide signals that, in addition to maintaining the GC by selecting GC B cells for survival and additional rounds of proliferation and mutation in the DZ, again influence GC B cell differentiation into memory B cells or plasma cells. GC-derived plasma cells and long-lived plasma cells produce the high affinity, class switched antibodies critical to pathogen clearance and long-term immunity; while different subpopulations of GC-derived memory B cells are able to rapidly differentiate into plasma cells or re-initiate the GC upon re-exposure to antigen.

Several Tfh-derived signals have been identified that can, through genetic deletion or antibody blockade, influence B cell differentiation. These include the cytokines IL-4 and IL-21 (Linterman et al., 2010; Weinstein et al., 2016) and receptors PD-1 and ICOS (Good-Jacobson et al., 2010; Liu et al., 2015). It is possible that differential expression of these factors is the mechanism by which the immune system tailors the B cell response to different antigens, but this has not been explored. BCR affinity for antigen is known to influence B cell fate choice, with higher affinity being linked to preferential plasma cell differentiation (Paus et al., 2006), but how or if an antigen can influence the cognate T cell partner or the signal it provides to B cells is not known.

Recent advances in understanding GC development and the cognate B/T interactions that drive them have benefited from model antigen systems in which B and T cells specific for the antigen can be identified and their activation and differentiation tracked over the course of the response. For example, we and others have transferred fluorescent ovalbumin (OVA)-specific T cells isolated from OTII mice and nitrophenyl hapten (NP)-specific B cells from B1-8 mice to non-fluorescent mice to track both cell types in the developing GC following immunization with NP-haptenated OVA (NPOVA) (Kerfoot et al., 2011; Shulman et al., 2014). Similar models based on other, almost always foreign antigens produce very similar outcomes. A model system based on an autoantigen may provide a tool with which to dissect the mechanisms by which the immune system itself controls differential outcomes, without relying on external blockade or deletion of candidate factors, yet the development of the autoimmune GC is under explored.

Myelin oligodendrocyte glycoprotein (MOG) is a well characterized autoantigen associated with anti-myelin autoimmunity of the central nervous system, both in human multiple sclerosis (MS) and the well-characterized animal model experimental autoimmune encephalomyelitis (EAE). In MS, anti-myelin B cells and antibodies show evidence of somatic hypermutation, indicating that they are GC-derived (Stern et al., 2014; von Büdingen et al., 2012). Currently, the most common way to induce MOG autoimmunity in C57Bl/6 mice is to immunize with the MOG_35-55_ peptide that corresponds to the CD4^+^ T cell epitope, a method that excludes B cell targeting of the MOG protein (Dang et al., 2015). However, we have shown that immunization with a larger peptide corresponding with the MOG-extracellular domain does indeed result in GC development incorporating anti-MOG B cells (Dang et al., 2015; Tesfagiorgis et al., 2017). Therefore, we assembled and developed the tools necessary to generate a MOG-based model antigen system analogous to the NPOVA system described above for investigation of differential B cell fate choice under the control of notably different antigens.

Here, we demonstrate that the GC develops very differently in response to MOG compared to the well characterized NPOVA system. In comparison to the NPOVA response, B cell fate choice in the MOG GC response was heavily biased against plasma cells. Further, while the MOG GC developed normally, it was not sustained and instead collapsed early, producing a large number of memory-phenotype cells. By manipulating the T cell pairing, we determined that, while plasma cell differentiation was largely independent of T cell influence, while class switch, GC maintenance, and differentiation into memory-phenotype cells were largely under the control of the T cell partner. By manipulating the antigen itself, we for the first time found the T cell affinity for antigen impacts B cell fate choice. Finally, we determined that memory phenotype cells derived from the MOG GC are not responsive to secondary challenge, and that this is intrinsic to the B cell and not due to education by the autoimmune T cell. To the best of our knowledge, this is the first example of unresponsive B cells derived directly from the GC.

## Methods/Materials

### Mice

C57Bl/6, 2D2 TCR-transgenic (Bettelli et al., 2003), SMARTA TCR-transgenic (4694; Tg(TcrLCMV)327Sdz/JDvsJ), and OTII TCR-transgenic mice (4194; Tg(TcraTcrb)425Cbn/J) were purchased from Jackson Laboratories, Bar Harbor, Maine. B1-8 mice (Maruyama et al., 2000) with a homozygous deletion of the J! locus (Chen et al., 1993) were a generous gift from Dr. Ann Haberman. IgH^MOG^ MOG-specific BCR knockin mice (Litzenburger et al., 1998) were received as a gift from Dr. H Wekerle. Mice expressing fluorescent proteins within all nucleated cells, either dsRed (RFP; 6051; Tg(CAG-DsRedpMST)1Nagy/J) under control of the “-Actin promoter or eGFP via the ubiquitin promoter (4353; Tg(UBCGFP)30Scha/J) were obtained from the Jackson Laboratory. Mice were housed in a specific pathogen-free barrier at West Valley Barrier. All animal protocols (2011-047) were approved by the Western University Animal Use Subcommittee.

### Antibodies for histology/flow cytometry

The following antibodies were purchased from BD Biosciences, Franklin Lakes, New Jersey: anti-Bcl6 A647 or v450 (K112-91), anti-CD138 BV421 or biotin (281-2), anti-CXCR5 APC (2G8), anti-CD19 BV711 (1D3), anti-CD4 v450 (RM4-5), anti-CD62L A700 (Mel14), anti-CD95 PE-Cy7 (Jo2), anti-IgG1 APC (A85-1), Streptavidin v450 or APC-Cy7, and anti-CD80 PE (16-10A1). The following antibodies were purchased from Thermo Fisher Scientific, Waltham, Massachusetts: anti-BrdU A647 (MoBU-1), anti-IgM A568 (polyclonal), anti-CXCR4 PE (2B11), Streptavidin A568, anti-Ki67 unconjugated. The following antibodies were purchased from eBioscience, Waltham, Massachusetts: anti-PD-1 biotin (RMP1-30), anti-CD38 PE or PE-Cy5 (90), anti-CD4 PE-Cy5 (RM4-5), anti-FoxP3 eF660 (FJK-16s), anti-IgD eF450 (11-26c), anti-IgG1 PerCP-eF710 (M1-14D12), Streptavidin APC, anti-ICOS biotin (C398.4A), and anti-PD-L2 biotin (TY25). The following antibodies were purchased from BioLegend, San Diego, California: anti-His Tag purified (J099B12), anti-PD-1 PE-Cy7 (RMP1-30), anti-rabbit IgG Dylight 649 (polyclonal), anti-CD4 A647 (RM4-5).

### Cloning of haMOG_tag_

The pET-32 mMOG_tag_ vector (Jain et al., 2016) was mutated by PCR using the following primers: 5’ TCTTCTTTTTCTCGCGTTTCTGGTTCTCCGTCTTCTGGTTTTGAAAACTTGTATTTCCAAGGACAGTTTCGCG 3’ and the reverse primer 5’ GCGAGAAAAAGAAGAACGGGTTTCGGTAACACGACGATATGCACCGGAGCCACCACCGGTAC 3’. The resulting vector was sequenced to confirm the insertion of the 13-35 neurofilament-M sequence and transformed into BL21 bacteria for expression.

### MOG production and purification

mMOG_tag_ and haMOG_tag_ proteins were produced and purified as previously described (Jain et al., 2016). The final equimolar concentrations were 5 mg/mL for mMOG_tag_ and 5.394 mg/mL for haMOG_tag_ with no detectable impurities as determined by SDS-PAGE.

### Adoptive transfer of B and T cells and immunization

Naïve antigen-specific T cells were isolated from RFP^+^ 2D2 and OTII mice and naïve antigen-specific B cells were isolated from GFP^+^ IgH^MOG^ and B1-8 J!^-/-^ mice as previously described (Kerfoot et al., 2011). Briefly, lymph nodes and spleens of RFP^+^ antigen-specific T cell and GFP^+^ antigen-specific B cell mice were dissociated and B and T cells were isolated using EasySep Negative selection Mouse B and T cell Enrichment Kits (StemCell Technologies, Vancouver, Canada). Unless otherwise stated, 5 # 10^5^ RFP^+^ T cells and either 1 x10^6^ GFP^+^ B1-8 J!^-/-^ or 5 # 10^6^ GFP^+^ IgH^MOG^ B cells (to account for the fact that only 20% are MOG-specific (Dang et al., 2015)) were transferred i.v into C57Bl/6 or SMARTA recipients 2 d prior to immunization. Mice were immunized in the footpad with equimolar amounts of the given antigen (125 µg mMOG_tag,_ 175 µg NPOVA, 125 µg NPMOG_tag_ (both at a 1:25 protein:NP ratio), 135 µg haMOG_tag_) in CFA. Unless otherwise stated, draining popliteal lymph nodes were harvested at the indicated time points for analysis. In experiments using BrdU, 1.5 mg of BrdU was injected i.p at the specified time points.

### Flow cytometry

Draining popliteal lymph nodes were harvested from mice for FACS analysis as previously described (Dang et al., 2015). Briefly, lymph node cell suspensions were blocked with an anti-Fc$ receptor, CD16/32 2.4G2 (BD biosciences), in PBS containing 2% FBS before further incubation with the indicated antibodies. Dead cells were excluded by staining with either the Fixable Viability Dye eFluor506 (eBioscience), propidium iodide (Thermoscientific), or 7-AAD (Biolegend). Flow cytometry was performed on a BD Immunocytometry Systems LSRII cytometer and analyzed with FlowJo software (Tree Star, Ashland, Oregon). For intracellular stains of FoxP3 or Bcl6, cells were fixed and permeabilized with Cytofix / Cytoperm solution (BD Bioscience) after cell surface staining. Fixed cells were then intracellularly stained for Bcl6 and FoxP3 at 4°C overnight. For BrdU staining, cells were fixed in 2% PFA then permeabilized in 0.1% Tween 20 for two nights at 4°C. The DNA within the fixed cells was degraded using DNase I (Sigma-Aldrich, St. Louis, Missouri) then stained with anti-BrdU antibody. Cell sorting was performed using a BD FACS ARIAIII where cells were sorted into 100% FBS.

### Immunofluorescent histology

Tissues were prepared for histology as previously described (Dang et al., 2015). Briefly, whole popliteal lymph nodes were fixed in periodate–lysine–paraformaldehyde (PLP), subsequently passed through sucrose gradients to protect from freezing artifacts and then frozen in OCT (TissueTek, Torrance, California) media. Serial cryostat sections (7 µm) were blocked in PBS containing 1% Bovine Serum Albumin, 0.1% Tween-20 and 10% rat serum before proceeding with staining. Sections were mounted with ProLong Gold Antifade Reagent (Invitrogen, Carlsbad, California). Tiled images of whole lymph node sections (20#) were imaged using DM5500B fluorescence microscope (Leica, Wetzlar, Germany).

### T cell proliferation assay

RFP^+^ OTII or 2D2 CD4^+^ T cells were enriched through negative selection as described above. Splenocytes of wild type C57Bl/6 mice were depleted of red blood cells using ACK lysis buffer (Thermo Fisher Scientific). The cells were then transferred into 10% FBS RPMI with L-glutamine (Thermo Fisher Scientific) supplemented with 1x penicillin/streptomycin (WISENT, Saint-Bruno, Canada). One million splenocytes were then added to individual wells of a sterile 48-well plate and were incubated with either 35 %g NP-OVA, 25 %g mMOG_tag_, or 27 %g haMOG_tag_ for one hour at 37 °C 5% CO2. OTII or 2D2 T cells were CFSE (Thermo Fisher Scientific) labelled as previously described (Jain et al., 2016) and 4 x 10^5^ T cells were added to the antigen loaded splenocytes. After three days of co-culture, CFSE labeling of antigen-specific T cells was analyzed by flow cytometry.

### Digital Droplet PCR (ddPCR)

Tfh and naïve T cells were sorted by flow and RNA was extracted from cells using a RNeasy Plus Micro Kit (QIAGEN, Hilden, Germany) and immediately converted into cDNA using a Superscript VILO cDNA Synthesis Kit (Invitrogen). ddPCR reactions were set up using ddPCR EvaGreen 2x Supermix (Bio-Rad, Hercules, California) and the following primers: IL-4 Sense – 5’ AGATGGATGTGCCAAACGTCCTCA 3’, IL-4 Antisense – 5’ AATATGCGAAGCACCTTGGAAGCC 3’, IL-10 Sense – 5’ GGTTGCCAAGCCTTATCGGA 3’, IL-10 Antisense – 5’ ACCTGCTCCACTGCCTTGCT 3’, IL-21 Sense – 5’ TGAAAGCCTGTGGAAGTGCAAACC 3’, IL-21 Antisense – 5’ AGCAGATTCATCACAGGACACCCA 3’, CD40L Sense – 5’ GTGAGGAGATGAGAAGGCAA 3’, CD40L Antisense – 5’ CACTGTAGAACGGATGCTGC 3’, ICOS Sense – 5’ TGACCCACCTCCTTTTCAAG 3’, ICOS Antisense – 5’ TTAGGGTCATGCACACTGGA 3’, PD-1 Sense – 5’ CGTCCCTCAGTCAAGAGGAG 3’, PD-1 Antisense – 5’ GTCCCTAGAAGTGCCCAACA 3’, CD28 Sense – 5’ TGACACTCAGGCTGCTGTTC 3’, CD28 Antisense – 5’ TTCCTTTGCGAGAAGGTTGT 3’, CTLA4 Sense – 5’ GCTTCCTAGATTACCCCTTCTGC 3’, CTLA4 Antisense – 5’ CGGGCATGGTTCTGGATCA 3’, FoxP3 Sense – 5’ CCCAGGAAAGACAGCAACCTT 3’, FoxP3 Antisense – 5’ TTCTCACAACCAGGCCACTTG 3’. ddPCR reactions were run on a QX200 Droplet Digital PCR System (Bio-Rad) and analyzed using Quantasoft software (Bio-Rad). Gene expression was normalized to the number of sorted cells and expressed as mRNA copies per cell.

### ELISpots and ELISA

96-well plates were coated overnight at 4°C with 0.5 µg NPOVA, NPMOG_tag_, or mMOG_tag_. Wells were blocked with 1% (wt/vol) BSA in PBS, then incubated with serial diluted bone marrow or lymph node cells at 37°C in 5% CO_2_. Spots were detected using a goat alkaline phosphatase-conjugated anti-mouse IgM or IgG antibody (MABTECH, Nacka Strand, Sweden) and 5-bromo-4-chloro-3-indolyl-phosphate substrate (Sigma-Aldrich) and counted under a Leica M80 dissection microscope. To detect circulating antibodies using an ELISA, 96-well plates were incubated with antigen and blocked with BSA as written above. Blood was extracted from mice using a cardiac puncture and spun at 4500 x g for 15 minutes. Serum plasma was extracted and incubated with the 96-well plate for one hour at room temperature. Plates were incubated with anti-IgM or IgG antibodies and then the alkaline phosphatase yellow (pNPP; Sigma-Aldrich) substrate. OD405 was measured using an Eon microplate spectrophotometer (BioTek, Winooski, Vermont).

### Image and statistical analyses

Histology images were analyzed using ImageJ software to quantify the density of B and T cells in germinal centers (Bcl6^+^ IgD^-^) and B cell follicles (IgD^+^ cells excluding five cells deep worth of the outermost perimeter of the B cell follicle near the capsule). PRISM software (GraphPad Software, La Jolla, California) was used to analyze FACs and histology data. For statistical comparisons, a students T-test was used for single comparisons and a one-way ANOVA followed by a T test with Bonferroni correction was used for multiple comparisons.

## Results

### Immunization with MOG autoantigen results in an atypical, unsustained GC response

In order to identify and track responding B and T cells throughout an immune response to two different antigens, GFP^+^ B cells (either NP-specific B1-8^+^ Jκ^-/-^ or MOG-specific IgH^MOG^) and RFP^+^ T cells (either OVA-specific OTII or MOG-specific 2D2) were isolated from mutant mice and transferred into wild type C57BL/6, non-fluorescent recipients (Figure 1A). Two days post transfer, mice were immunized in the footpad with the appropriate antigen (NPOVA for recipients of B1-8 B cells and OTII T cells, or mMOG_tag_ for recipients of IgH^MOG^ B cells and 2D2 T cells) in CFA. Lymph nodes were harvested for histological analysis 5d post immunization, representing the outcomes of early, pre-GC cognate interactions between responding B and T cells, or 10d post immunization, representing a mature GC time point.

**Figure 1:**
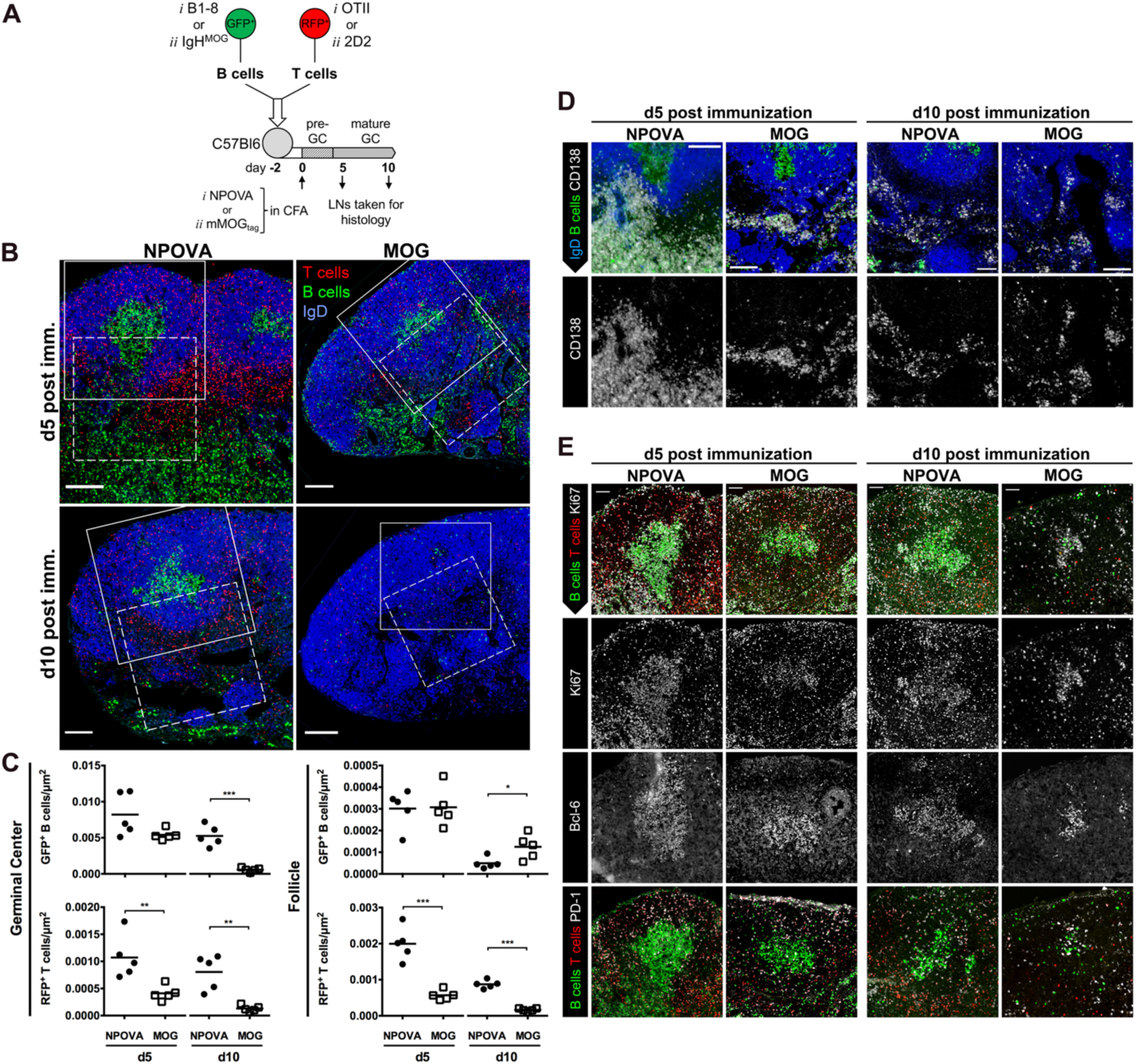
Differential GC development in the NPOVA and MOG model antigen systems. (**A**) Fluorescent B and CD4^+^ T cells specific for NPOVA or MOG were isolated and transferred into wild type, non-fluorescent recipients. Two days post-transfer, mice were immunized with either NPOVA or mMOG_tag_ in CFA in the footpad. Draining popliteal lymph nodes were harvested for histology d5 and d10 post immunization, representing the early and mature GCs. (**B**) Immunofluorescence of lymph nodes from NPOVA and mMOG_tag_ immunized mice to visualize RFP^+^ T cells and GFP^+^ B cells derived from transferred antigen-specific cells. Sections were also stained for IgD to outline B cell follicles. Scale bars represent 100 %m. (**C**) The density of GFP^+^ or RFP^+^ cells in the GC or follicle was quantified. Each data point represents the average value across one histological section for a single mouse. * p<0.05, **p<0.01, ***p<0.001. (**D**) Higher magnification of the regions of interest outlined by the dashed lines in panel B showing CD138 staining for plasma cells. (**E**) Higher magnification of the regions of interest outlined by the solid white line in panel B were further examined for Ki67, Bcl6, and PD-1 expression.

While virtually no transferred fluorescent cells could be observed in lymph nodes from unimmunized mice (Not Shown), large numbers of fluorescent B and T cells derived from the original transferred populations were readily evident at the 5d time point in both antigen systems (Figure 1B top, C). Consistent with our previous observations (Kerfoot et al., 2011) PD-1^+^ RFP^+^ Tfh cells were distributed throughout the follicle and GC in both model systems, although the density of RFP^+^ T cells was significantly lower in mMOG_tag_-immunized mice (Figure 1B, C, E).

Very large numbers of GFP^+^ CD138^+^ cells, representing the early short-lived plasmablast response, were evident outside of the follicles and within medullary cords of NPOVA- but not mMOG_tag_-immunized mice (Figure 1D). By 10d post immunization fewer, but equivalent numbers of plasma cells were within medullary cords in both model systems.

Within B cell follicles, dense clusters of GFP^+^ cells (Figure 1B, C) that were also IgD^lo^, Ki67^+^, and Bcl-6^+^ (Figure 1A, E) were evident in both systems 5d post immunization, indicating that early pre-GC B/T interactions were sufficient to drive GC B cell differentiation and establishment of a new GC. However, by 10d post immunization, the GC in the MOG antigen system had largely disappeared, while this time point corresponded with the full development of a mature and organized GC in the NPOVA system (Figure 1B bottom, C). Small clusters of Ki67^+^ and Bcl-6^+^ cells could still be observed in follicles of mMOG_tag_-immunized mice, however these were much smaller and less dense than those observed in NPOVA mice (Figure 1E). Instead, greater numbers of individual GFP^+^ cells were scattered throughout the follicle (Figure 1B, C). Very few individual GFP^+^ cells were evident in the follicle in the NP-OVA system, and virtually all remained confined in the GC.

### Preferential differentiation of B cells with a memory phenotype in response to MOG autoantigen

The developing GC response was analyzed by FACS in a separate, identical experiment. Consistent with our histological observations, the early CD19^int^ CD138^+^ plasma cell response was nearly absent in mMOG_tag_-immunized mice compared to a very large response in the NPOVA system (Figure 2A, B). This was true of both the GFP^+^ response derived from transferred, antigen specific B cells and the endogenous GFP^-^ response (Figure 2B bottom), confirming that this is a feature of the anti-MOG response.

**Figure 2:**
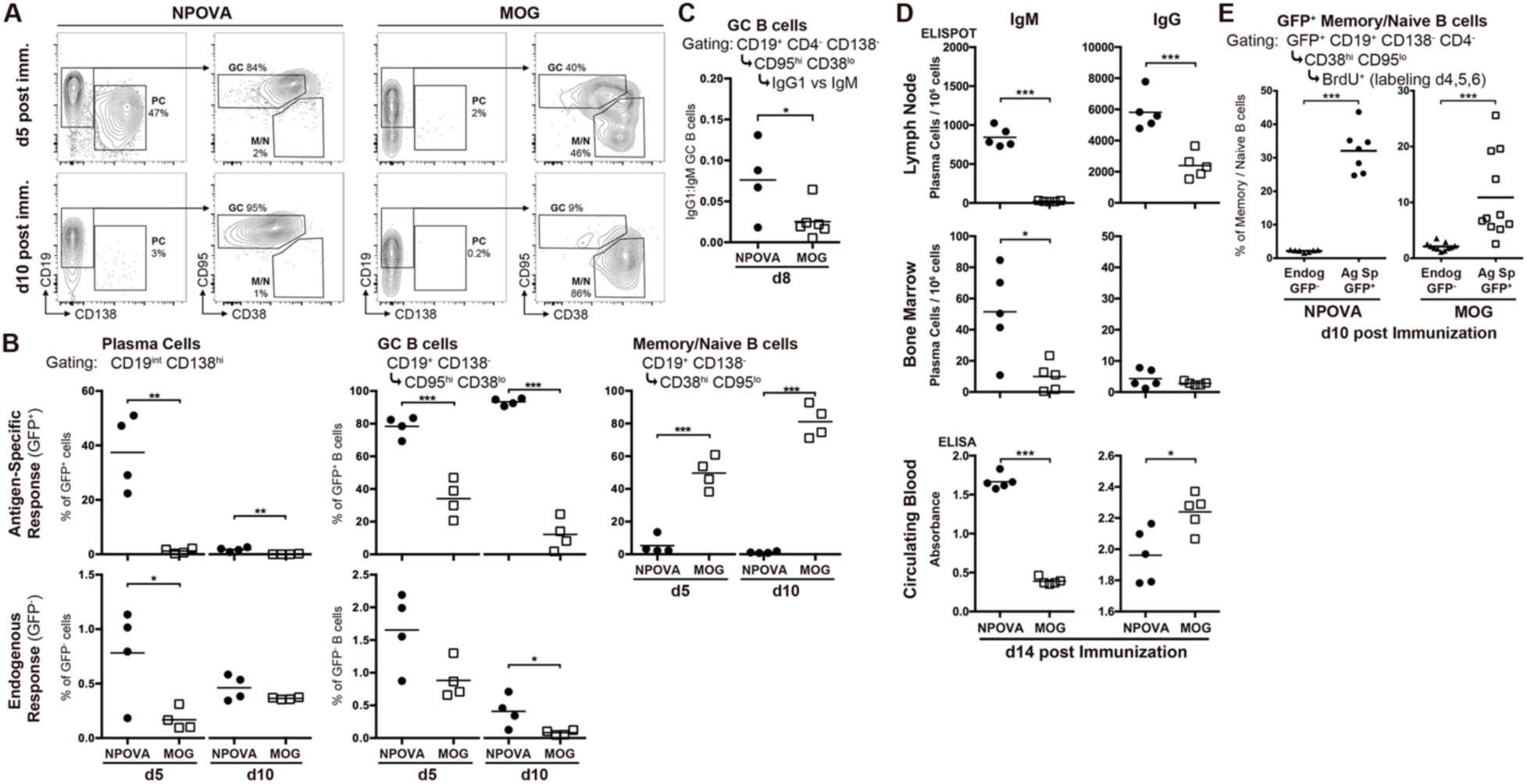
Early collapse of the MOG GC to a memory phenotype. Fluorescent B and CD4^+^ T cells specific for NPOVA or MOG were transferred into non-fluorescent C57Bl/6 mice that were then immunized with NPOVA or mMOG_tag_. Draining lymph nodes were harvested for analysis by FACS d5 and d10 post-immunization. (**A**) Representative gating of GFP^+^ cells for plasma cells (PC), GC B cells, and memory/naive B cells (M/N). (**B**) Quantification from panel A showing size of each cell subset (as defined in panel A, gating shown above each plot) derived from the transferred GFP^+^ B cells (top row) or from endogenous GFP^-^ cells (bottom row). Data is expressed as the percentage of all GFP^+^ cells for Plasma cells, or percentage of all GFP^+^ B cells (CD19^+^ CD138^-^) for GC and Memory/Naïve B cells. One representative of two separate experiments is shown. (**C**) The ratio of IgG1 expressing cells over IgM expressing GC B cells d8 post-immunization is shown. (**D**) C57Bl/6 mice were immunized with either NPOVA or mMOG_tag_ in CFA. d14 post-immunization, draining popliteal lymph nodes and bone marrow were taken for ELISpot analysis of NP- or MOG-specific IgM or IgG. Blood serum from the same mice was assayed by ELISA for circulating anti-NP or -MOG IgM or IgG antibodies. (**E**) Fluorescent antigen-specific B and T cells were transferred into non-fluorescent SMARTA recipient mice and immunized with NPOVA or mMOG_tag_. Mice were injected i.p. with BrdU d4, d5, and d6 post-immunization and BrdU incorporation in the GFP^+^ or GFP^-^ memory/naïve B cell populations was assessed by FACS d10 post-immunization. Each graph represents a separate experiment and the data points in the mMOG graph were pooled from two separate experiments. In all graphs, each data point represents an individual mouse. * p<0.05, **p<0.01, ***p<0.001.

While antigen-specific GFP^+^ CD95^hi^ CD38^lo^ GC B cells were evident in both the NPOVA and MOG systems at the d5 time point, they made up a significantly smaller proportion of the total GFP^+^ B cell population in the MOG system (Figure 2A, B), and most dramatically at the 10d time point, consistent with the collapse of the GC response observed by histology. A similar collapse of the endogenous, GFP^-^ GC was also observed in mMOG_tag_ immunized mice (Figure 2B bottom). The proportional loss of GFP^+^ antigen-specific GC B cells and plasma cells in the MOG response was offset by a large increase in the proportion of CD38^hi^ CD95^lo^ cells, a phenotype shared by naïve and memory B cells (Figure 2B, top right).

Evaluation of class switch in the GC B cell population 8d post immunization, prior to complete collapse of the MOG GC, revealed that the ratio of IgG1 to IgM-expressing GC B cells was significantly higher in NPOVA-immunized mice (Figure 2C). Nevertheless, and despite the bias against plasma cell development (Figure 2A, B), mMOG_tag_-immunized mice were still capable of mounting an antigen-specific antibody response, albeit smaller than the response observed in response to NPOVA. Indeed, by ELISpot the number of anti-MOG IgM and IgG producing cells was significantly lower in lymph nodes 14d post immunization compared to anti-NP producing cells (Figure 2D top). Similar analysis of bone marrow revealed a reduction in anti-MOG IgM, but not IgG-producing cells (Figure 2D middle). This was reflected by reduced levels of circulating anti-MOG compared to anti-NP IgM but not IgG, as measured by ELISA of serum from the same mice (Figure 2D bottom).

### Antigen-specific GFP^+^ CD38^hi^ CD95^lo^ B cells are antigen experienced

To confirm that the GFP^+^ CD38^hi^ CD95^lo^ B cells observed above derive from previously activated and proliferating cells, BrdU was injected 4, 5, and 6d post immunization to label proliferating cells. On day 10-post immunization, lymph nodes were harvested for FACS analysis. In this way, only cells that were actively proliferating during the labeling period (note that only a proportion of actively proliferating cells would be labeled, due to the short half-life of free BrdU in mice), but had then become quiescent would retain BrdU labeling (Weisel et al., 2016). Indeed, neither non-proliferating endogenous GFP^-^ CD38^hi^ CD95^lo^ follicular B cells (Figure 2E), nor proliferative GFP^+^ CD95^hi^ CD38^lo^ GC B cells (not shown) stained with BrdU. In contrast, a proportion of GFP^+^ CD38^hi^ CD95^lo^ memory/naïve B cells were BrdU^+^ in both model systems, confirming that they derived from previously activated cells.

Combined, our histology and FACS findings demonstrate that, compared to a standard well-studied model foreign antigen, the B cell response to MOG protein produces a short-lived GC response with relatively little class switch and reduced plasma cell differentiation. Instead, the GC response dissolves early to produce a large number of memory-phenotype, non-proliferating cells distributed throughout the follicle.

### T cells partially control the outcome of the GC response to MOG

To begin to decipher the role for the cognate T cell partner in instructing differential B cell fate choice and the failure of GC maintenance in the MOG vs NPOVA systems, we took advantage of the modular nature of the hapten antigen system to place NP-specific B1-8^+^ Jκ^-/-^ B cells under control of either OVA-specific OTII T cells (NPOVA) or MOG-specific 2D2 T cells (NPMOG).

Fluorescent NP-specific B cells were transferred to non-fluorescent recipients expressing an irrelevant transgenic TCR (SMARTA) in order to limit the endogenous T cell response. Either OVA or MOG-specific T cells were transferred at the same time. Recipients were immunized 2d later with the appropriate antigen and lymph nodes were harvested 5 or 10d post immunization for analysis by FACS or, in a separate experiment, histology.

Similar to the response to MOG observed above (Figure 2B), 5d post immunization short-lived plasmablasts made up a smaller proportion of the NP-specific GFP^+^ response under control of MOG-specific T cells compared to OVA specific T cells (Figure 3A) although the difference was not as extreme and, unlike in the response to MOG, plasma cell numbers had fully recovered by d10. A large GC was evident 5d post-immunization by FACS (Figure 3A) and histology (Figure 3B, C) in both systems, indicating that OVA and MOG-specific T cells are capable of supporting the early formation of a GC. However, by 10d post immunization there was evidence that the NPMOG GC had begun to collapse, as GC B cells made up a smaller proportion of the total antigen-specific population compared to the NPOVA response (Figure 3A), and GCs were less dense (Figure 3B, C). This was balanced by a significant increase in the proportion of antigen-specific B cells with a memory/naïve CD38^hi^ CD95^lo^ phenotype (Figure 3A). Further, class switch on GC B cells was also significantly reduced under the control of MOG-specific T cells (Figure 3D). Therefore, ongoing maintenance rather than initiation of the GC, as well as class switch, are in part controlled by the T cell partner of the cognate B/T pairing.

**Figure 3:**
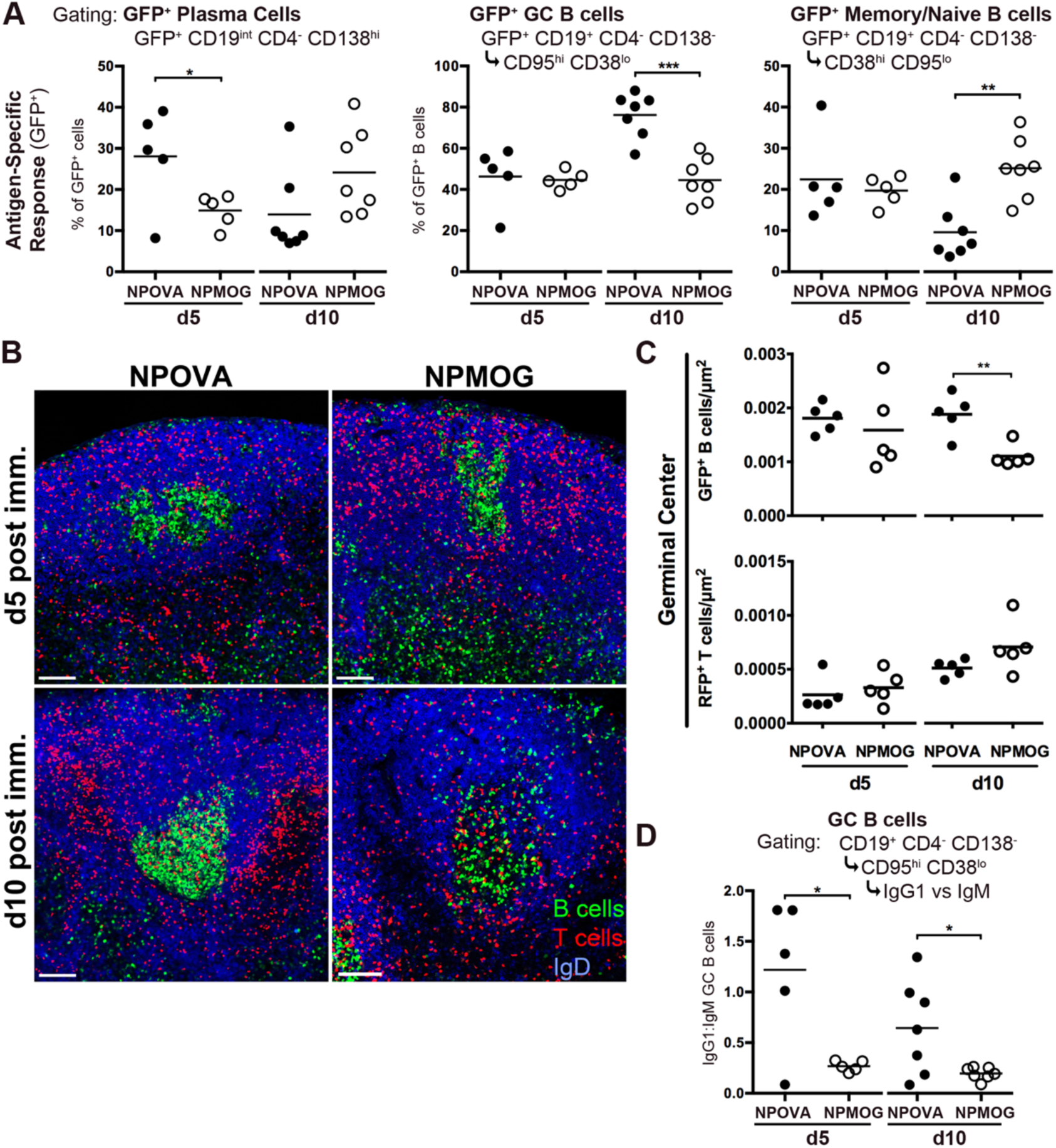
MOG-specific T cells induce early GC collapse to a memory phenotype. Fluorescent NP-specific B cells and either OVA or MOG-specific CD4^+^ T cells were transferred into non-fluorescent SMARTA recipients that were then immunized with either NPOVA or NPMOG. Draining popliteal lymph nodes were harvested for analysis by FACS or, in a separate experiment, histology d5 and d10 post immunization. (**A**) The size of the given cell subset is shown as a percentage of all GFP^+^ cells (Plasma cells) or all GFP^+^ B cells (GC B cells and Memory/Naïve B cells). The d5 and d10 time points were assessed in separate experiments, data shown is representative of 2 to 3 individual experiments. (**B**) Representative histological sections from NPOVA or NPMOG-immunized mice to visualize NP-specific GFP^+^ B cells and either RFP^+^ OVA-specific or MOG-specific T cells, respectively. Sections were stained for IgD to outline B cell follicles. (**C**) The density of GFP^+^ or RFP^+^ cells in the GC was quantified from histological images. (**D**) The ratio of IgG1-over IgM-expressing GC B cells d5 and d10 post-immunization was determined in a separate FACS experiment. Each data point represents an individual mouse. * p<0.05, **p<0.01, ***p<0.001.

### Low T cell antigen affinity limits the MOG GC response

A common feature of autoimmune TCRs, including TCRs that recognize the MOG_35-55_ peptide, is that they tend to bind peptide:MHC with relatively low affinity (Deng and Mariuzza, 2007; Ramadan et al., 2016). Many are also polyreactive – meaning that they recognize more than one specific peptide. Indeed, analysis of the MOG_35-55_-specific 2D2 TCR revealed that it also recognizes a second peptide derived from the Neurofilament-M protein (NF-M_18-30_) (Krishnamoorthy et al., 2009), and in fact binds NF-M_18-30_ with higher affinity than it does MOG_35-55_ (Rosenthal et al., 2012). We took advantage of polyreactivity of the 2D2 TCR to determine if TCR affinity for antigen influences B cell fate choice and maintenance of the GC response by generating a modified mMOG_tag_ antigen that incorporates the NF-M_18-30_ epitope (Figure 4A – referred to as “high affinity” or haMOG_tag_). Initial validation experiments were performed to confirm processing and presentation of haMOG_tag_ to T cells. Isolated, CFSE-labeled OTII or 2D2 T cells were cultured with splenocytes loaded with NPOVA, mMOG_tag_, or haMOG_tag_. 2D2 T cell proliferation to haMOG_tag_ was intermediate, between that of OTII cells in response to NPOVA and 2D2 cells in response to mMOG_tag_ (Figure 4B).

**Figure 4:**
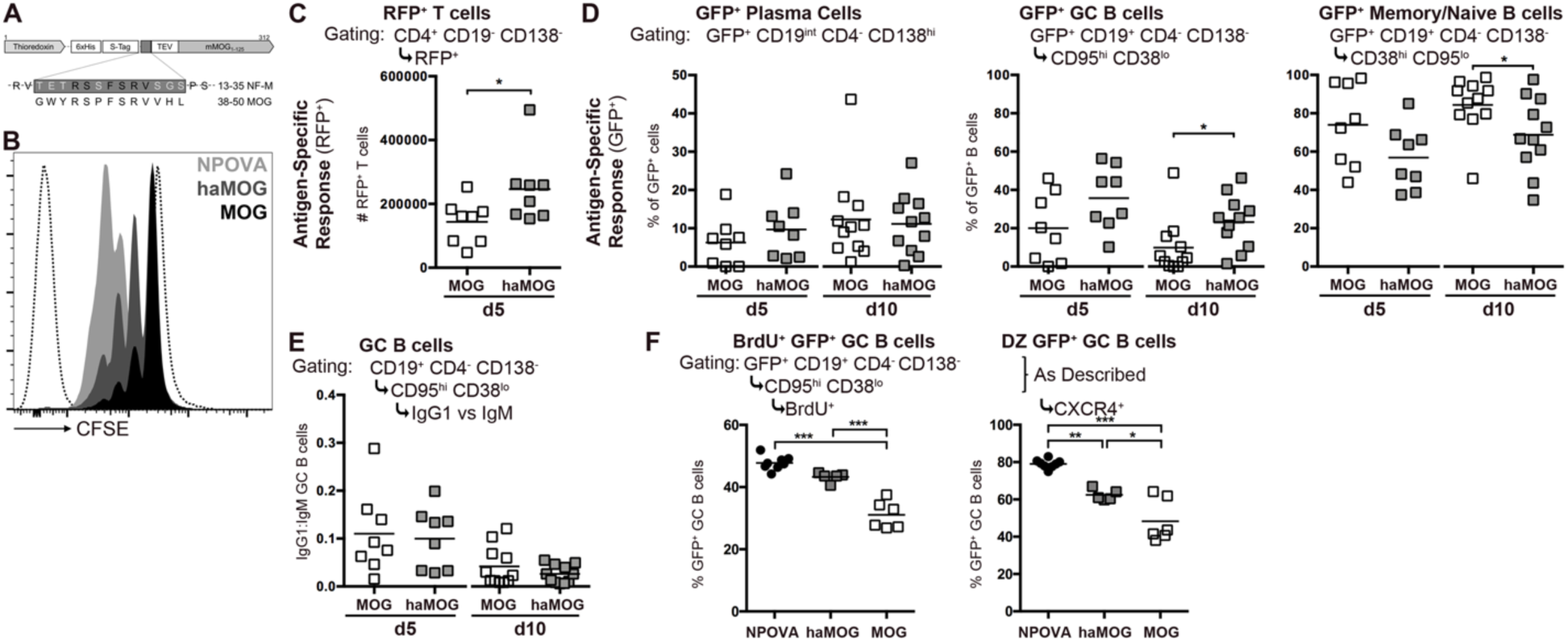
Increasing T cell antigen affinity partly rescues the MOG GC from early collapse. (**A**) A schematic of the haMOG_tag_ Ag showing the insertion of amino acids 13-35 from neurofilament-M (with sequence comparison to the MOG_35-55_ peptide) (**B**) *In vitro* proliferation assay measuring CFSE-dilution of labeled OVA-specific T cells cultured for 3d with NPOVA-loaded splenocytes, or labeled 2D2 T cells cultured with mMOG_tag_, or haMOG_tag_-loaded splenocytes. A representative (of 3 separate experiments) histogram for each condition is shown. The dashed lines represent unlabeled (left) and fully CFSE-labelled (right) OTII T cells. (**C-E**) Fluorescent MOG-specific B and T cells were transferred into non-fluorescent SMARTA recipients that were then immunized with mMOG_tag_ or haMOG_tag_. Draining popliteal lymph nodes were harvested for analysis by FACS d5 and d10 post-immunization. The d5 and d10 time points were assessed in separate experiments, data shown is the combination of two separate experiments. (**C**) The absolute number of RFP^+^ T cells is shown d5 post-immunization. (**D**) The size of the given cell subset at both 5 and 10d post immunization is shown as a percentage of all GFP^+^ cells (Plasma cells) or all GFP^+^ B cells (GC B cells and Memory/Naïve B cells). (**E**) The ratio of IgG1-over IgM-expressing cells was determined for GC B cells. (**F**) Fluorescent Ag-specific B and T cells were transferred into non-fluorescent SMARTA recipients that were then immunized with NPOVA, mMOG_tag_, or haMOG_tag_. Mice were injected i.p. with BrdU 7d post immunization, and draining popliteal and inguinal lymph nodes were harvested for analysis by FACS 12hrs later. The percentage of GFP^+^ GC B cells that are BrdU^+^ (left) or CXCR4^+^ (right) is shown. Each data point represents an individual mouse. * p<0.05, **p<0.01, ***p<0.001.

Having validated the haMOG_tag_ antigen, fluorescent MOG-specific B and T cells were transferred to SMARTA recipients which were then immunized with either mMOG_tag_ or haMOG_tag_. Lymph nodes were harvested 5 or 10d later for analysis by FACS. Greater numbers of RFP^+^ 2D2 T cells were recovered from haMOG_tag_ immunized compared to MOG_tag_ immunized mice (Figure 4C), confirming that, as in our *in vitro* assay, haMOG_tag_ induces greater T cell proliferation *in vivo*. No differences in plasma cell differentiation were observed at either time point (Figure 4D). However, consistent with the hypothesis that the TCR affinity of the T cell partner in the cognate pair influences GC maintenance vs B cell differentiation, partial recovery of the GC with a corresponding decrease in the proportion of memory-phenotype B cells was observed 10d post immunization with haMOG_tag_. In contrast to our observations where NP-specific B cells were placed under control of two different T cells (Figure 3), T cells responding to haMOG_tag_ did not affect class switch in the GC (Figure 4E), suggesting that these outcomes are controlled separately or that they represent a gradient of potential outcomes influenced by different levels of T cell activation and signal production.

In the cyclic reentry model of the GC response (Victora and Nussenzweig, 2012), GC B cells undergo repeated rounds of proliferation and somatic hypermutation, largely in the dark zone (DZ), followed by migration to the light zone (LZ) to receive survival and differentiation signals, predominantly from Tfh cells. We hypothesized that the collapse of the MOG GC was due to the inability of Tfh cells to drive LZ B cells to maintain GC status and reenter the DZ, instead resulting in differentiation to a memory phenotype. To test this, proliferation of GC B cells was analyzed by BrdU uptake, along with the expression of CXCR4 as a marker of DZ GC B cells. Consistent with our hypothesis, BrdU labeling of GC B cells was significantly higher in the NPOVA system compared to either the mMOG_tag_ or haMOG_tag_-immunized mice, and more GC B cells were of the CXCR4^+^ DZ phenotype, while haMOG_tag_-induced GCs were intermediate (Figure 4F).

### Levels of T cell activation do not explain the differential B cell response between the different model systems

In an attempt to understand the underlying mechanism behind the differential outcome of the GC response in the different model antigen systems, antigen-specific Tfh cells (CXCR5^+^ PD-1^hi^ RFP^+^) were FACS sorted from lymph nodes of mice 10d post immunization with NPOVA, mMOG_tag_, or haMOG_tag_ (Figure 5A, B). mRNA was isolated for quantitative digital droplet PCR analysis of the expression of proteins with a known role in providing T cell help and differentiation signals to GC B cells. Surprisingly, little difference was observed in expression levels of the canonical Tfh cytokines IL-4 and IL-21 (the small difference in IL-4 expression was not consistent across experiments) nor the expression of IL-10 (Figure 5C, top). Neither were their differences in the expression of the surface receptors CD40L, ICOS, PD-1, CD28 and CTLA-4 (Figure 5C, middle). Equivalent surface expression of ICOS and PD-1 by antigen-specific Tfh cells was confirmed in a separate experiment by FACS (Figure 5D). Interestingly, the master regulator of regulatory T cells, FoxP3, was expressed at significantly higher levels by Tfh cells from haMOG_tag_-immunized mice (Figure 5C bottom), a finding confirmed by FACS (Figure 5D). The significance of this observation is not clear, as an increased ratio of T follicular regulatory cells would seem to counter the larger GC response in haMOG_tag_ vs MOG_tag_-immunized mice. Nevertheless, this finding was consistent across three separate ddPCR and FACS experiments.

**Figure 5:**
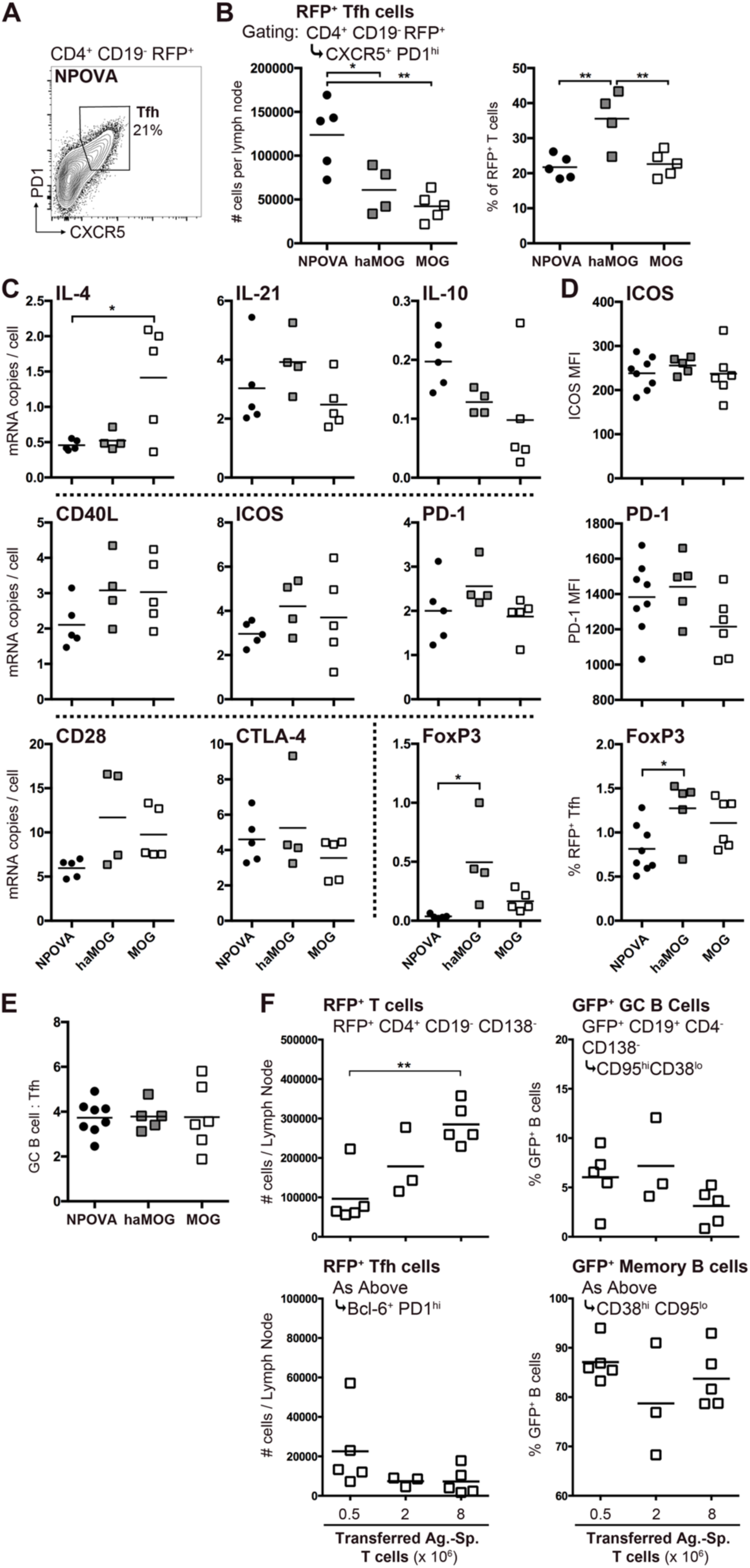
Tfh cell phenotype is not altered by antigen. (**A-C**) RFP^+^ antigen-specific T cells were transferred along with non-fluorescent antigen-specific B cells into non-fluorescent SMARTA recipient mice that were then immunized with NPOVA, mMOG_tag_, or haMOG_tag_ in CFA. Draining popliteal and inguinal lymph nodes were harvested 10d later and Tfh cells (CD4^+^ CD19^-^ RFP^+^ CXCR5^+^ PD-1^hi^) were FACS sorted for subsequent analysis of gene expression by digital droplet PCR. One representative of two independent experiments is shown. (**A**) An example of gating for CXCR5^+^ PD-1^hi^ Tfh cells is shown. (**B**) The absolute number of Tfh cells per lymph node is shown (left panel), along with size of the Tfh population as a percentage of total RFP^+^ T cells (right panel). (**C**) Digital droplet PCR analysis of mRNA levels (copies per cell) for the listed gene. (**D-E**) Fluorescent antigen-specific B and T cells were transferred into non-fluorescent SMARTA recipients that were then immunized with NPOVA, mMOG_tag_, or haMOG_tag_. Draining popliteal and inguinal lymph nodes were harvested 8d post immunization for analysis by FACS. (**D**) Mean fluorescence intensity (MFI) for ICOS and PD-1 on RFP^+^ CXCR5^+^ PD-1^hi^ Tfh cells (top two panels) and the percent of Tfh cells (Bcl6^+^ PD-1^hi^) that were FoxP3^+^ (bottom panel) are shown. (**E**) Ratio of GC B cells to Tfh cells in the different antigen systems. (**F**) Fluorescent MOG-specific B cells and different numbers of MOG-specific T cells (0.5, 2, or 8 x 10^6^ 2D2 T cells) were transferred into non-fluorescent SMARTA recipients that were then immunized with mMOG_tag_. Draining popliteal lymph nodes were harvested 10d post immunization for FACS analysis. The absolute number of RFP^+^ T cells per lymph node (top left panel) and RFP^+^ Tfh cells per lymph node is shown (bottom left panel). The percentage of the GFP^+^ B cells with a GC B cell (top right panel) or memory B cell (bottom right panel) phenotype is shown. Each data point represents an individual mouse. * p<0.05, **p<0.01.

We consistently observed that the absolute number of Tfh cells was greater in the NPOVA vs MOG systems (Figure 5B, and also reflected in Figure 1C) and that haMOG_tag_ immunization produced intermediate numbers of Tfh cells (Figure 5B, and also reflected in Figure 4C). This resulted in the GC B cell:Tfh cell ratio remaining the same across model antigen systems (one example presented in Figure 5E). To determine if the size of the GC response was simply linked to the size of the T cell response to a given antigen, different numbers of 2D2 T cells were transferred along with equal numbers of MOG-specific B cells into SMARTA recipient mice. While immunization with mMOG_tag_ resulted in a significantly larger antigen-specific T cell response in mice that received more cells, there was no similar increase in the number of Tfh cells, nor was there an alteration in the GC response (Figure 5F).

### MOG-induced memory B cells are not responsive to secondary challenge

The primary function of memory B cells is to respond to secondary immune challenge (Weisel and Shlomchik, 2017). To determine if CD38^hi^ CD95^lo^ memory phenotype B cells generated from the MOG GC are responsive to antigen challenge, we performed an experiment that isolates the primary and secondary responses within the same mouse (Figure 6A). After transfer of fluorescent, antigen-specific B and T cells, SMARTA recipients were immunized in the left footpad only and 34d later, the same mice were immunized in the right footpad. Left and right draining lymph nodes were analyzed separately by FACS 5d post secondary challenge. Control mice immunized with NPOVA in CFA in the left footpad but “challenged” with adjuvant alone showed an ongoing (but small in absolute terms – data not shown) GFP^+^ GC response in the left but not right draining lymph nodes (Figure 6B middle), confirming the lymphatic separation of the two sides. As expected, memory phenotype cells made up the vast majority of GFP^+^ cells on the right side, confirming that memory cells generated in the primary GC properly circulate and home to lymphatic tissues (Figure 6B bottom). As expected, secondary challenge with NPOVA resulted in generation of short lived plasmablasts (Figure 6B top) and initiation of a GC response on the right, but not the left side. This contrasted starkly with the challenge response in mMOG_tag_-immunized mice. Consistent with previous observations, the primary GC response on the left side in mMOG_tag_-immunized mice had disappeared, along with evidence of plasma cells at the 39d time point, leaving GFP^+^ cells with exclusively a CD38^hi^ CD95^lo^ phenotype. Despite the clear presence of memory-phenotype GFP^+^ cells in the right lymph node, secondary challenge with mMOG_tag_ antigen did not produce a new GC response or plasma cells.

**Figure 6:**
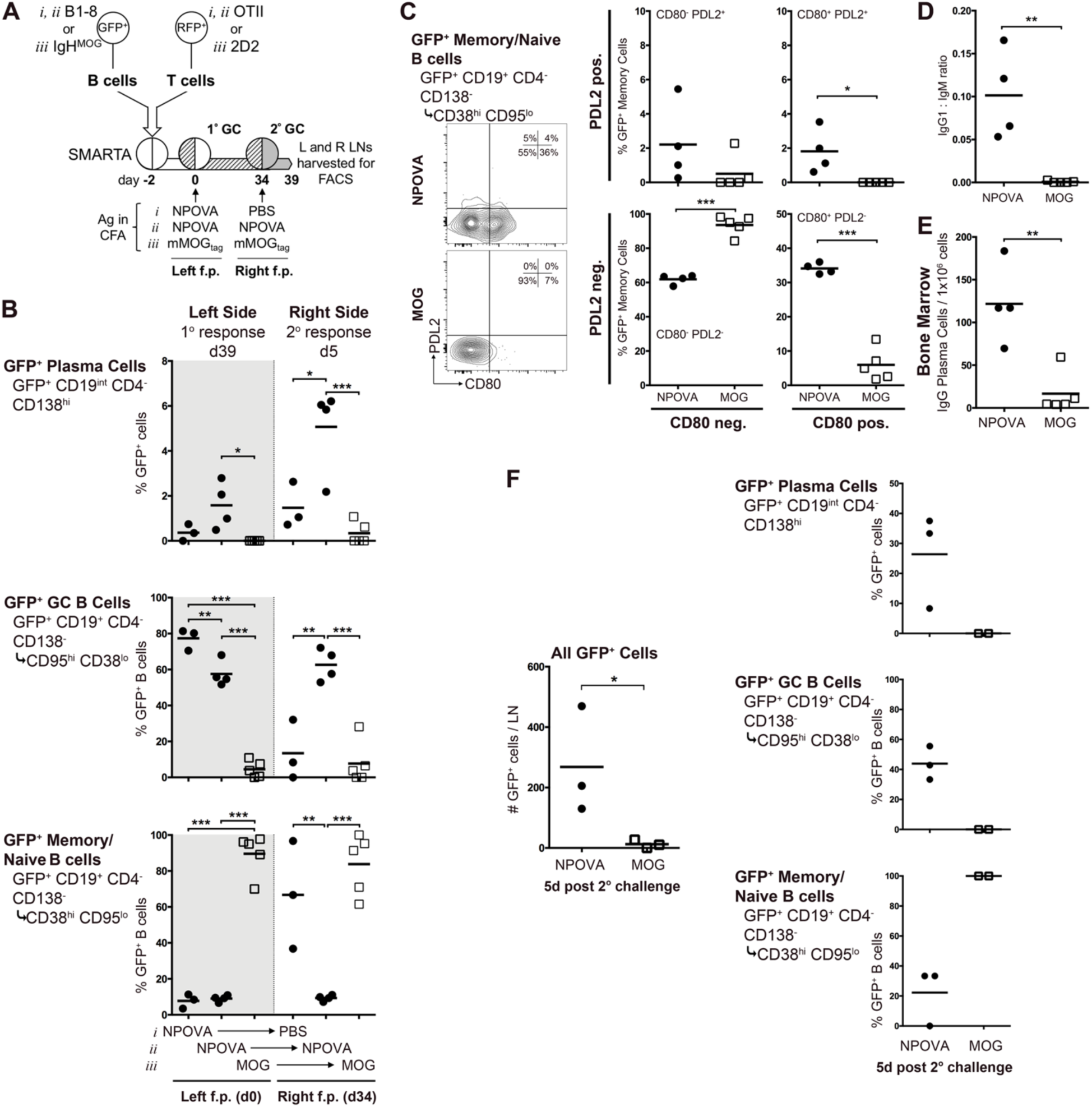
Memory B cells produced by the MOG GC response are unresponsive to secondary challenge. (**A**) Fluorescent antigen-specific B and T cells were transferred into non-fluorescent SMARTA recipients that were then immunized in their left footpad with either NPOVA or mMOG_tag_ in CFA. Thirty-four days post-immunization, mice were immunized in their right footpad with NPOVA, PBS, or mMOG_tag_ in CFA in the right footpad, as shown. (**B**) The primary response in the left draining popliteal and inguinal lymph nodes and secondary response in the right lymph nodes were analyzed separately by FACS 5d post challenge. The size of the given antigen-specific subsets as a percentage of the total GFP^+^ cells (Plasma cells) or GFP^+^ B cells (GC and Memory/Naïve B cells) is shown for the left and right sides separately. (**C**) Representative staining and quantification for CD80 and PD-L2 on NPOVA and MOG GFP^+^ memory/naïve B cell subsets. (**D**) The ratio of IgG1 expressing cells over IgM expressing cells amongst GFP^+^ memory/naïve B cells is shown. (**E**) At the same time, bone marrow was harvested for ELISpot quantification of NP- or MOG-specific IgG producing plasma cells. (**F**) Fluorescent antigen-specific B and T cells were transferred into non-fluorescent SMARTA recipients that were then immunized with NPOVA or mMOG_tag_ in CFA. Draining popliteal and inguinal lymph nodes were harvested 10d post immunization and CD19^+^ CD4^-^ CD138^-^ CD38^hi^ CD95^lo^ GFP^+^ memory/naïve B cells were sorted. 7.5 x 10^3^ cells were transferred along with 5 x 10^5^ naïve T cells specific for the appropriate antigen into new non-fluorescent SMARTA recipient mice. These were immunized with NPOVA or mMOG_tag_ and 5d later draining popliteal and inguinal lymph nodes were analyzed by flow cytometry. The absolute number of GFP^+^ cells per lymph node is shown (left) and then broken down by subset on the right. Each data point represents an individual mouse. * p<0.05, **p<0.01, ***p<0.001.

Recently, subsets of memory B cells have been identified based on differential expression of PD-L2 and CD80 (Tomayko et al., 2010). Double negative memory cells are associated with the establishment of a new GC (Zuccarino-Catania et al., 2014). Nevertheless, CD38^hi^ CD95^lo^ GFP^+^ B cells in mMOG_tag_-immunized mice were almost entirely double negative, while a significant proportion of memory cells in NPOVA immunized mice expressed PD-L2 and/or CD80 (Figure 6C). Class switch remained reduced on GFP^+^ memory cells in the MOG system compared to the NPOVA system (Figure 6D), and significantly fewer IgG-producing long-lived plasma cells were recovered from the bone marrow (Figure 6E).

In the above experiment, it is possible that the presence of Treg cells generated in the primary response to MOG inhibited the subsequent secondary response. To eliminate this possibility, GFP^+^ antigen specific CD38^hi^ CD95^lo^ memory phenotype cells were FACS sorted from mMOG_tag_ or NPOVA immunized mice and equal numbers were transferred to new SMARTA recipients along with naive T cells specific for the relevant antigen. Following secondary challenge, small numbers of GFP^+^ NP-specific cells were recovered, the majority of which were plasma cells or GC B cells (Figure 6F). In contrast, MOG-specific cells were either completely undetectable or exclusively of the CD38^hi^ CD95^lo^ phenotype, indicating that they had not responded to secondary challenge.

To determine if the unresponsiveness of MOG-specific memory B cells was due to education from MOG-specific T cells, an experiment was performed to determine if MOG-specific T cells could educate NP-specific B cells to be similarly unresponsive. After transfer of NP-specific B cells along with the appropriate OVA or MOG-specific T cells, recipient mice were immunized with NPOVA or NPMOG in the left footpad only. 32d later, mice received naive T cells specific for the reciprocal antigen and were then challenged with that antigen in the right footpad 2d later (Figure 7A). Left and right draining lymph nodes were analyzed separately by FACS 5d post secondary challenge. Analysis of the primary response in the left lymph node revealed that, as at d10 (Figure 3), the NP-specific B cell response under control of MOG-specific T cells was heavily biased to memory-phenotype cells at the expense of GC B cells (Figure 7B). The presence of IgG-producing long-lived plasma cells in the bone marrow was also reduced (Figure 7C). In contrast, and as opposed to the MOG-specific B cells in the previous experiment (Figure 6C), there was no difference in the proportion of CD80 PDL2 double negative memory NP-specific B cells under the control of either T cell (Figure 7D), nor was there a defect in class switch of memory cells (Figure 7E). Also, analysis of the right lymph node clearly demonstrated that NP-specific B cells educated by MOG-specific T cells in the primary response were able to respond to secondary challenge (Figure 7B).

**Figure 7:**
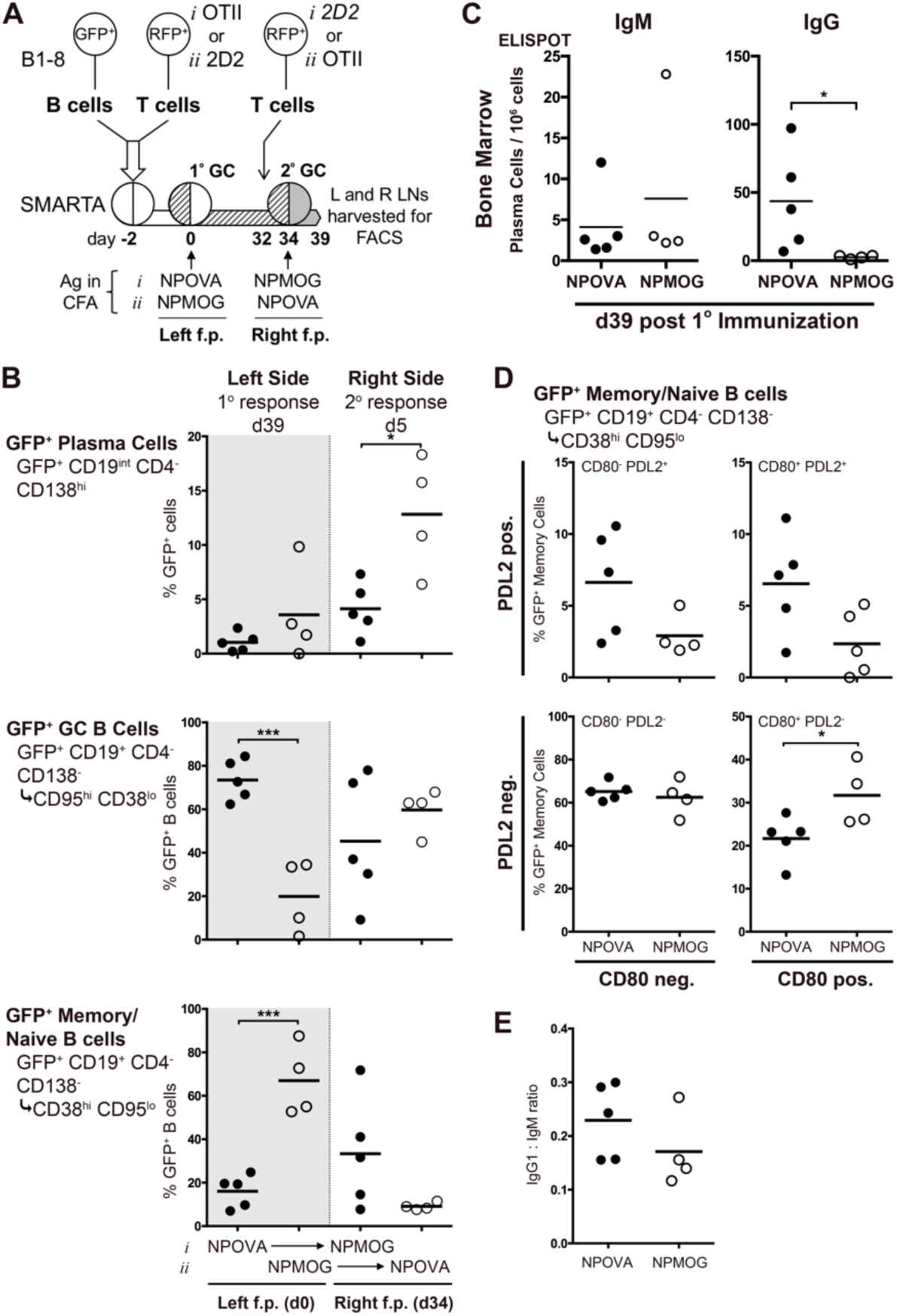
Autoimmune T cells do not induce unresponsiveness in MOG-specific B cells during the GC response. (**A**) Fluorescent NP-specific B cells and OVA or MOG-specific T cells were transferred into non-fluorescent SMARTA recipients that were then immunized in their left footpad with either NPOVA or NPMOG. Thirty-two days post-immunization, naïve T cells of the reciprocal specificity were transferred to these recipient mice, as shown, followed two days later by immunization with that antigen in the right footpad. (**B**) The primary response in the left draining popliteal and inguinal lymph nodes and secondary response in the right lymph nodes were analyzed separately by FACS 5d post challenge. The size of the given antigen-specific subsets as a percentage of the total GFP^+^ cells (Plasma cells) or GFP^+^ B cells (GC and Memory/Naïve B cells) is shown for the left and right sides separately. (**C**) At the same time, bone marrow was harvested for ELISpot quantification of NP-specific IgM or IgG producing plasma cells (the antigen used to coat plates was based on the primary immunogen). (**D**) Memory/naïve phenotype GFP^+^ B cells were analyzed for expression of CD80 and PD-L2. (**E**) The ratio of IgG1 over IgM expressing cells amongst GFP^+^ memory/naïve B cells is shown. Each data point represents an individual mouse. * p<0.05, **p<0.01, ***p<0.001.

## Discussion

Here, we use manipulatable antigen model systems as a novel approach to investigate how the immune system controls B cell fate choice and differentiation to produce different GC outcomes tailored to the specific antigen. The response to NPOVA and other NP haptenated proteins is well characterized (Shlomchik and Weisel, 2012; Weisel et al., 2016), and in many ways is considered to represent the default response to a foreign antigen. We and others have shown that the anti-NP GC consistently forms 4-5d after exposure to antigen, peaks ∼2 wks post exposure, and remains active for several weeks after that (Kerfoot et al., 2011; Zuccarino-Catania et al., 2014). We show here that, while the GC response to MOG develops with similar kinetics, it is not sustained and instead dissociates early. This should not be interpreted as failed GC response, however, as it still produces measurable levels of circulating, class switched anti-MOG antibody. Further, subcutaneous immunization with MOG protein is a well-established method to induce the anti-myelin autoimmune model experimental autoimmune encephalomyelitis (EAE). In our hands, mice immunized with mMOG_tag_ develop a robust disease with evidence that GC-derived anti-MOG B cells contribute to both disease severity and chronic disease course (Dang et al., 2015; Tesfagiorgis et al., 2017). Therefore, although short-lived, the MOG GC is productive.

The GC responses is sustained by interactions between GC B cells and Tfh cells, predominantly in the LZ of the GC. The outcome of these interactions can select B cells to maintain their GC status and cycle back into the DZ for additional rounds of cell division, mutation, and return to the LZ for selection (Mesin et al., 2016). Alternatively, GC B cells can be driven to memory or plasma cells fates (Suan et al., 2017). The first major finding of our study is that, in the MOG GC response, early failure of the GC is due to preferential differentiation to a memory phenotype at the expense of maintaining the GC. Indeed, within the small GC B cell population in the collapsing MOG response there is a clear bias to a LZ phenotype, suggesting that cells are not being selected to return to the DZ for proliferation. A similar bias to memory cell differentiation is seen for B cells defective in CXCR4, which is required for proper DZ B cell homing (Bannard et al., 2013). By histology, this manifests as a small, less-organized GC with a large number of individual GFP^+^ cells distributed throughout the follicle. In the GC response to a foreign antigen, memory B cell differentiation has been shown to occur predominantly in the early stages, shortly after GC formation, with plasma cell differentiation preferentially occurring later in the response (Weisel et al., 2016). Therefore, it is possible that the early dissolution of the MOG GC to generate memory B cells represents an extreme acceleration of this same process.

It is clear from our observations that the status of the cognate T cell partner strongly influences the dichotomy between GC maintenance and memory B cell differentiation, along with class switch. Indeed, MOG-reactive T cells induced a similar GC outcome when paired with NP-specific B cells and enhanced T cell activation via high affinity antigen partly rescued the MOG GC from collapse and reduced memory B cell differentiation. In this case, class switch was not impacted, suggesting that there is a gradient to the GC parameters that are influenced by T cell status. Interestingly, while BCR affinity has previously been linked to plasma cell differentiation (Kräutler et al., 2017; Paus et al., 2006) (see below), this is the first report that we are aware of that demonstrates that TCR affinity for antigen can impact B cell fate choice.

It is not clear what signals the cognate T cell partners use to drive differential GC maintenance vs memory B cell differentiation in the two model systems. Previously identified T cell signals that influence GC formation and memory differentiation include ICOS and PD-1 (Good-Jacobson et al., 2010; Liu et al., 2015). Tfh-produced cytokines, IL-21, IL4, and IL-10 have also been shown to be required for proper GC development (Laidlaw et al., 2017; Linterman et al., 2010; Weinstein et al., 2016). Nevertheless, we did not find evidence that these are differentially expressed by Tfh cells in the NPOVA and MOG systems. Therefore, the immune system may employ other signals to modulate GC outcome in response to different antigens. The size of the Tfh cell pool itself may be one of these “signals”, as we consistently observed a direct correlation between the number of Tfh and GC B cells in our different model systems. An attempt to modulate this by increasing the total T cell response was not successful, suggesting that other factors limit the size of the Tfh cell niche in an antigen-dependent way. Indeed, maintenance of the PD-1^hi^ phenotype on Tfh cells is dependent on their ongoing cognate interactions with B cells (Baumjohann et al., 2011; Kerfoot et al., 2011). Therefore, it is difficult to separate cause and effect with regards to the GC B cell:Tfh cell ratio.

While the balance between GC maintenance and memory B cell differentiation, along with class switch, were heavily influenced by the status of the cognate T cell partner, plasma cell differentiation and memory B cell unresponsiveness were not. Plasma cell differentiation has been linked to BCR affinity (Kräutler et al., 2017; Paus et al., 2006). Further, plasma cells preferentially differentiate later in the GC response compared to memory cells (Weisel et al., 2016). It is possible that the MOG GC doesn’t last long enough to produce BCRs with sufficiently high affinity to promote plasma cell differentiation. The accumulation of somatic mutations in anti-MOG B cells and BCR affinity for antigen will need to be explored in future studies. However, this would not explain the almost complete absence of early, short lived plasmablasts that typically derive from pre-GC interactions. The starting affinity for antigen in the Ig-heavy chain knockin (IgH^MOG^) B cells is clearly sufficient to allow for B cell activation to proliferate and initiate the GC. Additional investigation will be required to determine if (potentially) low BCR affinity accounts for reduced plasma cell formation, or if the few (but productive) plasma cells that do form in the MOG GC response represent clones that attained a threshold affinity that allowed for their differentiation.

Finally, we believe that this is the first demonstration of unresponsive memory B cells derived from a GC response, although they may be related to so-called “atypical” CD27^-^ CD21^-^ memory B cells identified in humans (Weisel and Shlomchik, 2017), and reported to be enriched in autoimmune conditions including MS (Claes et al., 2016). These B cells have been reported to be partly anergic (defined as having reduced BCR signaling capacity), however their role in driving or limiting inflammation is not well understood. It is not yet clear if anergy is the mechanism behind the memory B cell unresponsiveness in our MOG system. As with plasma cell differentiation, this non-responsiveness was not influenced by the status of the T cell partner as MOG-specific T cells did not educate NP-specific memory B cells to become non-responsive. Moreover, fresh naïve T cells were not able to rescue the anti-MOG memory B cells from their non-responsive state. The CD80^-^ PD-L2^-^ double negative status of these memory-phenotype cells was also not the result of T cell education and may be linked to their non-responsiveness, although double negative memory cells have previously been associated with preferential GC formation following secondary challenge (Zuccarino-Catania et al., 2014). Further study will be required to determine if this induced non-responsiveness is the result of tolerance mechanisms resulting from previous exposure to endogenous MOG antigen. Importantly, MOG-specific B cells are not initially unresponsive, as they generate a GC in the primary response, but only become unresponsive post-activation.

In conclusion, we show here that different antigens can drive GC responses with very different outcomes. Further, we identify GC maintenance vs memory B cell differentiation as a fate decision dichotomy that is regulated independently from plasma cell differentiation, and that the status of the cognate T cell partner heavily influences the former, but not the latter. Finally, we show that B cells can be induced during the GC response to be unresponsive to secondary challenge. Our findings have implications both for our fundamental understanding of how B cell fate choice is regulated in the GC response, and for our understanding of how autoimmune B cells participate in autoimmune responses, and anti-myelin responses in particular.

## Acknowledgements

The authors would like to thank the veterinarians and animal care staff at the West Valley Barrier Facility for their excellent husbandry of our experimental animals. RWJ is the recipient of an endMS Doctoral Studentship from the Multiple Sclerosis Society of Canada (MSSOC). KAP is the recipient of a endMS Post-Doctoral Fellowship from MSSOC. YT is supported by an Ontario Graduate Scholarship.

## Author Contributions

RWJ performed the bulk of the experiments and contributed to conceptualization and writing. KAP, YT, and HBC contributed experiments, and ER participated in image analysis. SMK provided supervision and contributed to writing.

